# Representational Competition of Spatially and Temporally Overlapped Target and Distractor

**DOI:** 10.1101/2025.10.15.682617

**Authors:** Changhao Xiong, Ke Bo, Lihan Cui, Nathan M. Petro, Andreas Keil, Mingzhou Ding

**Affiliations:** J. Crayton Pruitt Family Department of Biomedical Engineering, University of Florida, Gainesville, FL 32611; Department of Psychological and Brain Science, Dartmouth College, Hanover, NH 03755; Boys Town National Research Hospital, Boys Town, NE 68010; Department of Psychology, University of Florida, Gainesville, FL 32611

**Keywords:** Attention, distractor, fMRI, competition, multivariate pattern analysis

## Abstract

Representational competition occurs when a task-relevant target stimulus and a distractor overlap in space and time. Given limited neural resources, it is expected that stronger representations of the distractor will result in weaker representations of the target, leading to poorer behavioral performance. We tested this hypothesis by recording fMRI while participants (n = 27) viewed a compound stimulus consisting of randomly moving dots (the target) superimposed on IAPS affective pictures (the distractor). Each trial lasted ∼12 seconds, during which the moving dots and the IAPS pictures were flickered at 4.29 Hz and 6 Hz, respectively. The task was to detect and report brief episodes of coherent motion in the moving dots. Depending on the emotion category of the IAPS pictures, the trials were classified as pleasant, neutral and unpleasant. Focusing on three ROIs: middle temporal cortex (MT), ventral visual cortex (VVC), as well as the primary visual cortex (V1), we performed MVPA analysis to decode the distractor categories in each ROI, and correlated the decoding accuracy, taken to index the strength of distractor representation, with the accuracy in detecting the episodes of coherent motion. The following results were found: (1) the decoding accuracy was above chance level in all ROIs, and (2) in MT and VVC but not in V1, the higher the decoding accuracy, the worse the behavioral performance. These results suggested that distractor information was represented in V1 as well as in the two motion-processing areas, and in the motion-processing areas, stronger representations of the distractor led to poorer ability to process attended information, leading to worse behavioral performance. The hypothesis was thus supported and trade-offs in the fidelity of stimulus representations prompted by neural competition demonstrated.

## Introduction

At any given moment, the richness and complexity of visual information entering the retina exceeds the capability of the capacity-limited visual system. Selective visual attention is essential for prioritizing relevant information (e.g., targets) while filtering out irrelevant information (e.g., distractors). When targets and distractors appear in different spatial locations, the human brain uses visual spatial selective attention mechanisms to resolve competition (Moran & Desimone, 1985; Reynolds et al., 1999). According to the biased competition model (Desimone & Duncan, 1995), cortical representations of targets interact with that of distractors via reciprocal inhibition (Reynolds & Desimone, 2003), with stronger representations of targets leading to weaker representations of distractors. In the case of spatial selective attention, the dorsal attention network, comprised of intraparietal sulcus and frontal eye fields, is thought to play an essential role in this process by sending top-down biasing signals that boost neural activity at the task-relevant location and concurrently down-weight task-irrelevant locations (Corbetta & Shulman, 2002). The effectiveness of this location-based mechanism is however weakened when targets and distractors are spatially proximate (Driver & Baylis, 1989; Eriksen & Eriksen, 1974). Visual spatial attention becomes ineffective when the targets and the distractor fully overlap in space and time. Other mechanisms are required for implementing selection, including object-based, and feature-based attention.

Visual objects are characterized along multiple feature dimensions such as color, shape, or motion. Attention can be directed to any one of these features or combination of features. In this case, the unattended features become task-irrelevant information. Neurons specialized for different visual features are distributed across distinct brain regions: color-sensitive neurons in areas V2, V4, and associated ventral stream regions (Conway, 2009; Lueck et al., 1989; Zeki, 1978); motion-sensitive neurons in the middle temporal area (MT/MST) and related dorsal stream structures (Treue & Trujillo, 1999); shape-sensitive neurons in the lateral occipital complex (LOC; (Grill-Spector et al., 1998; Kourtzi & Kanwisher, 2000; Malach et al., 1995); and orientation-sensitive neurons in primary visual cortex V1 (Hubel & Wiesel, 1968; Ringach et al., 2002). This anatomical segregation has given rise to the hypothesis that feature-based selection mechanisms is accomplished by selectively modulating the neural gain in feature-specific cortical areas (Boylan et al., 2023; McMains et al., 2007). In support of this hypothesis, it has been demonstrated that attending to a particular feature facilitates neural responses in the corresponding feature-selective cortical regions and pathways while suppressing neural activity that processes competing features (Andersen et al., 2009; McAdams & Maunsell, 2000). Such a push-pull mechanism may also apply to more complex viewing conditions with multiple features and naturalistic stimuli—the hypothesis examined in the present study.

Emotionally engaging (pleasant and unpleasant) pictures have been shown to act as strong competitiors for limited capacity, interfering with a wide range of primary, concurrent tasks (Bradley, 2009; Müller et al., 2008; Wieser & Keil, 2011). Recent work has shown that the emotional content of naturalistic scenes is represented in visual cortical areas (Bo et al., 2021). More specifically, when divided into three broad emotion categories: pleasant, neutral and unpleasant, the emotional category of the stimulus can be decoded from the spatial fMRI patterns evoked by the stimulus. When task-irrelevant images (distractors) spatially and temporally overlap with a task-relevant stimulus, it is therefore expected that within-area competition will play an important role in determining task performance. To test this idea, we recorded fMRI data from human subjects performing a motion detection task. The stimulus was a cloud of randomly moving dots (the target) superimposed on emotional pictures from the International Affective Picture System (IAPS; (Lang et al., 1997) serving as the distractor. The target and the distractor were flickered at two different frequencies (4.29 Hz vs 6 Hz) for an extended duration of ∼12 seconds. The participants were asked to focus on the randomly moving dots, ignore the IAPS pictures, and report the number of times the dots moved coherently. We extracted neural representations of distractor processing from the fMRI data and assessed (1) whether the distractor was represented in task-related brain regions and (2) how the representation of the distractor interfered with task performance.

## Materials and Methods

### Participants

The experimental protocol was approved by the Institutional Review Board of the University of Florida. A total of 30 undergraduate students from the University of Florida provided written informed consent and participated in the study. All participants were screened for MRI contraindications, including ferromagnetic implants, claustrophobia, and personal or familial history of epilepsy or photic seizures. Female participants also underwent a pregnancy test prior to participation. Three participants were excluded due to excessive movement during data acquisition. The fMRI data from the remaining n=27 participants (18 women, 9 men, mean age = 19.2 ±1.1 years) were analyzed and reported here.

#### Stimuli

The stimulus consisted of a random-dot kinematogram (RDK) superimposed on affective images selected from the International Affective Picture System (IAPS) database. The RDK consisted of 175 yellow dots, randomly distributed within a circular aperture centered on the screen, with each dot subtending less than 0.5 degrees of visual angle. The IAPS images were broadly divided into three emotional categories: pleasant, neutral, and unpleasant. These images were matched in overall composition, rated complexity, and file size to minimize potential confounds across categories. Stimuli were presented on a 30-inch MR-compatible LCD monitor positioned approximately 230 cm from the participant’s head outside the bore of the MRI scanner and viewed through a mirror mounted on the head coil. A central white fixation point remained visible throughout the experiment.

#### Procedure

See Figure 1(A) for a schematic illustration of the experimental task. On each trial, the subject was presented a compound stimulus array consisting of the RDK superimposed on the IAPS pictures. The moving dots and the IAPS pictures were flickered on and off at 4.29 Hz and 6 Hz, respectively. For each 4.29 Hz flicker cycle, the moving dots were displayed for 100ms, which was followed by a 133ms off period. Similarly, for each 6 Hz flicker cycle, the IAPS background picture was shown for 100ms and followed by a 66.7ms off period. During each on-off cycle, the moving dots in the RDK were randomly displaced by 0.3 degrees of visual angle in either random directions or one coherent direction. Coherent motion instances lasted for four on-off cycles (933 ms) and appeared once in 39 trials (13 trials per emotion condition) or twice in 4 trials. The remaining 41 trials contained no instances of coherent motion. Each trial lasted 11.667 seconds (50 moving dots cycles and 70 IAPS background picture cycles). The coherent motion instances occurred in the interval between 2.3 seconds and 10.4 seconds post stimulus array onset. The participant was asked to fixate the center white dot during the trial, monitor the motion coherence of the random dots, and report the number of coherent motion instances at the end of the trial. Both the number of coherent motion instances and the underlying emotion category of IAPS image were randomized on each trial. A total of 42 IAPS pictures were equally divided into three content categories based on valence: pleasant (erotic couples), neutral (workplace people), or unpleasant (bodily mutilation). For each trial, only IAPS pictures from one emotion category were presented.

**Figure 1.**
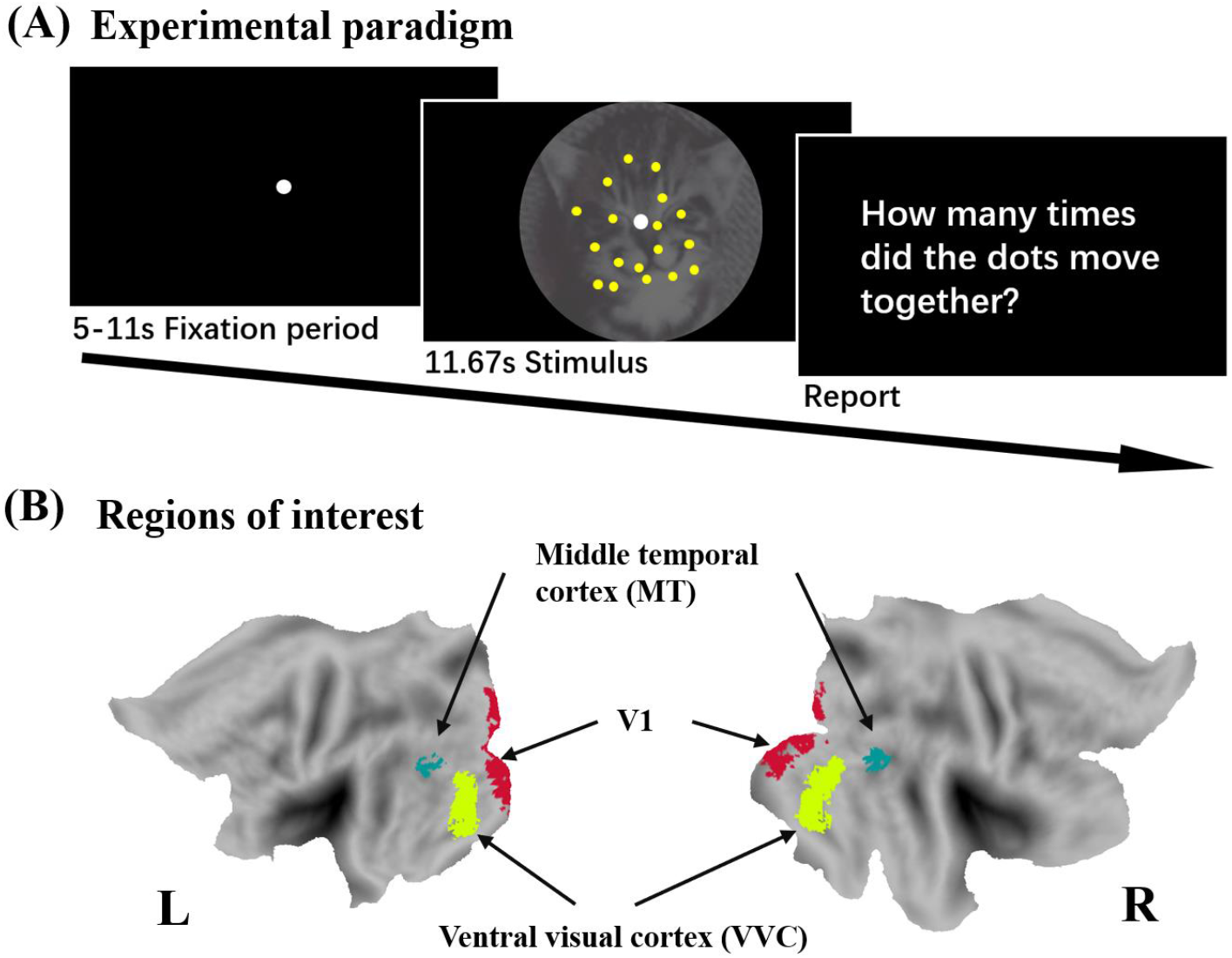
Paradigm and regions of interest. (A) The motion detection task paradigm. Randomly moving dots were superimposed on the IAPS pictures. Participants fixated the center white dot and reported the number of times the dots moved coherently (zero, one, or two) at the end of the trial (∼12 s in duration). (B) Regions of interest (ROIs): middle temporal cortex (MT), ventral visual cortex (VVC), and primary visual cortex (V1).

Depending on the emotion category of the IAPS pictures used, the trials are referred to as pleasant, neutral, and unpleasant trials, respectively. Each picture was used twice during the experiment. Consequently, there was a total of 84 trials: 28 pleasant trials, 28 unpleasant trials, and 28 neutral trials. According to this design, the moving dots were the target to be attended, whereas the IAPS pictures were the distractor to be ignored and suppressed.

#### fMRI data collection and preprocessing

The magnetic resonance imaging (MRI) data were acquired on a 3T Philips Achieva scanner using a gradient-echo echo-planar imaging (EPI) sequence with the following parameters: echo time (TE) = 30 ms, repetition time (TR) = 1.98 s, flip angle = 80°, 36 slices, field of view (FOV) = 224 mm, voxel size = 3.5 × 3.5 × 3.5 mm, and matrix size of 64 × 64. The first four functional scans were discarded to allow for scanner stabilization. Slices were collected in ascending order, aligned in parallel to the anterior-posterior commissure plane. Structural scans were obtained using a T1-weighted magnetization-prepared rapid gradient-echo (MP-RAGE) sequence with the following parameters: TE = 3.1 ms, TR = 8.1 ms, flip angle = 8°, 36 slices, FOV = 240 mm, voxel size = 1 × 1 × 1 mm, and matrix size = 240 × 240. EEG data were simultaneously acquired with the fMRI data but not considered here.

MRI data preprocessing was conducted using SPM aided by the ArtifactRepair toolbox (Mazaika et al., 2009) from the Center for Interdisciplinary Brain Sciences Research at Stanford Medical Center. All scans were realigned to the first scan in the sequence. Spatial smoothing was applied using a Gaussian kernel with a full width at half maximum of 4 mm. Movement correction was achieved through an algorithm in the ArtifactRepair toolbox designed to detect and correct significant residual errors in voxel translation due to participant motion. Large BOLD signal spikes caused by motion, defined as deviations exceeding 6% from the mean, were identified and interpolated using adjacent time points. After artifact correction, images were normalized and registered to the standard SPM template in Montreal Neurological Institute (MNI) space, with functional volumes resampled to a spatial resolution of 3 × 3 × 3 mm. Any remaining head motion was regressed out for each voxel across the entire session. Low-frequency drifts in the BOLD signal were removed using a 1/128 Hz high-pass filter and global signal normalization was performed by dividing each voxel by the mean value across slices.

#### ROI selection

Given that the main interest of this study was to examine how the distractor affected the processing of the motion stimulus, we focused on three regions of interest (ROIs, see Figure 1B): primary visual cortex (V1), the middle temporal (MT) cortex, and the ventral visual cortex (VVC). MT and VVC are known for their role in visual motion processing. In particular, the MT region’s essential role in processing motion and dynamic visual stimuli, including flickering stimuli, is well established (Born & Bradley, 2005; Newsome & Pare, 1988; Salzman et al., 1992). Regarding VVC, although it is primarily viewed as the region where object recognition and form perception take place, accumulating evidence suggests that VVC also plays an important role in motion processing. Patients with circumscribed lesions in the ventral visual cortex, which includes V4 and ventral occipital region (VO), exhibit severe impairments in central motion perception, rendering them unable to detect motion in the central visual field. Parahippocampal cortex (PHC), traditionally known for its scene recognition and spatial navigation functions, has also been implicated in motion processing, especially in distinguishing a moving object from its background (Aminoff et al., 2013). In the present study, V4, VO, and PHC were combined to form the VVC ROI. V1 was chosen as a ROI because it serves as the first cortical stage in the visual hierarchy where motion information is initially processed and relayed to higher-order motion areas such as MT and VVC (Lu & Sperling, 2001; Movshon & Newsome, 1996; Pascual-Leone & Walsh, 2001). Previous work has shown that different categories of IAPS images evoke distinct neural responses in these visual areas (Bo et al., 2021). As distractors here, we expected that they would compete for neural representations with the motion stimuli, which were the attended target.

The voxels for each of the three ROIs were selected based on a probabilistic brain atlas of the retinotopic visual cortex (Wang et al., 2015) and thresholded at 15/21 subjects—that is, for a given ROI, we retained only those voxels that were labelled as belonging to the ROI in ≥ 15 of the 21 atlas participants (probability ≥ 0.71). This conservative cutoff was chosen to maximize anatomical consistency across individuals.

#### fMRI data analysis

### Univariate analysis

For every run, the artifact-corrected BOLD signals were converted to percent-signal change by subtracting and then dividing by the run-wise mean. For each trial, we segmented the time series from 0 to 8 TR with 0 TR denoting the trial onset, and applied baseline correction.

### Multivariate analysis

In addition to univariate analysis, we also applied an MVPA decoding approach to assess the multivariate representation of the distractor in each of the ROIs as a function of time (i.e., TR). To enhance robustness, the data from two neighboring TRs were combined. For decoding, the linear support vector machine (SVM) method as implemented in the LibSVM package ((Chang & Lin, 2011) was used in conjunction with a leave-one-trial-out cross-validation method. More specifically, for each participant, and for a pair of emotional categories such as pleasant vs neutral or unpleasant vs neutral, we decoded pleasant distractors vs neutral distractors or unpleasant distractors vs neutral distractors. In each iteration of the cross-validation process, one trial was set aside as the test set, while the remaining trials were used to train the classifier. This process was repeated 56 times and the average of the 56 decoding accuracies on the test data was reported as the classification accuracy at each TR for the subject (Bae & Luck, 2019; Haxby et al., 2014; Zhang et al., 2024). Across participants, above-chance decoding accuracy is taken as evidence of distractor processing in the ROI, with higher decoding accuracy indicating stronger distractor representation.

## Results

Functional MRI data were recorded from participants who attended moving dots superimposed in IAPS pictures and detected instances of coherence motion while ignoring the distracting influence of the IAPS images. Each trial lasted ∼12 seconds during which the moving dots (RDK) were flickered at 4.29 Hz and the IAPS pictures flickered at 6 Hz. The participants reported the number of instances of coherent motion in the moving dots at the end of the trial (possible answers were 0, 1 or 2).

### Behavioral analysis

As shown in Table 1, the overall detection accuracy across the three trial types was 59.93% ± 2.55%, with that for pleasant, neutral, and unpleasant trials being 58.83% ± 2.72%, 59.32% ± 2.77%, and 61.65% ± 3.13%, respectively. A one-way ANOVA found no significant difference in behavioral performance between the three types of trials (F2, 44 = 0.273, p = 0.762). The task performance being similar across three types of distractors, two emotional and one neutral, may stem from the challenging nature of the motion detection task, as evidenced by the low overall accuracy (∼56%). In previous studies it has been shown that in a challenging task environment, more arousing distractors tend to have similar influences on task performance as neutral and less arousing distractors (Minamoto et al., 2015).

**Table 1.** Behavioral results. For pleasant, neutral, and unpleasant conditions, no significant difference was observed on task performance between trial types.

| Behavioral performance |  | Main effect of Content |
| --- | --- | --- |
| Overall | 59.93% ± 2.55% | $F_{2,44} = 0.273$<br>$p = 0.76$<br>$\eta^2 = 0.01$ |
| Pleasant | 58.83% ± 2.72% | |
| Neutral | 59.32% ± 2.77% | |
| Unpleasant | 61.65% ± 3.13% | |

### fMRI analysis: Univariate

Figure 2 plots univariate BOLD activation as functions of TRs for the three types of trials in MT, VVC, and V1. In all three ROIs, we observed significant BOLD activation evoked by the compound stimulus, the TR at which this occurred was denoted by an asterisk. Here statistical significance was defined as p<0.05 corrected for multiple comparison by the false discovery rate (FDR). Whereas the univariate analysis clearly demonstrated the involvement of the three ROIs in stimulus processing, a drawback is that it cannot separate the contribution due to the target from that due to the distractor.

**Figure 2.**
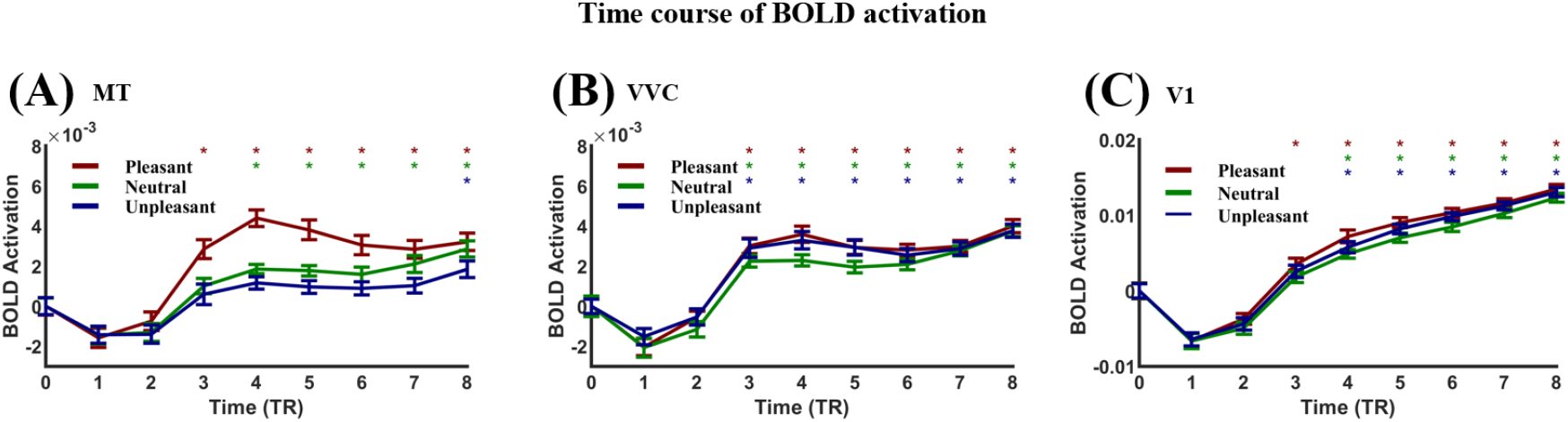
Univariate BOLD activation as function of TR in (A) MT, (B) VVC and (C) V1. The TRs at which the activation is significant are marked by an asterisk (p<0.05, FDR).

### fMRI analysis: Multivariate

To overcome the limitation of the univariate analysis and better characterize the competitive interaction between target and distractor representations, we implemented a multivariate decoding approach. Specifically, we took advantage of the distinct neural patterns associated with different types of emotional distractors (e.g., pleasant vs neutral), used the decoding accuracy to index the strength of distractor processing, and tested how the strength of the distractor neural representation influenced target detection performance. Our hypothesis was that the stronger the distractor representation, the weaker the target representations, the worse the behavioral performance.

#### Decoding accuracy time series

We combined the data from neighboring two TRs to increase the robustness of decoding. Figure 3 presents the temporal evolution of distractor decoding accuracy for pleasant vs. neutral (blue) and unpleasant vs. neutral (yellow) conditions in the three ROIs. In MT and VVC, decoding accuracy follows a similar trajectory, becoming above chance at TR 2-3 (4 to 6 seconds after stimulus onset), peaking at around TR 4-5, while in V1, the decoding accuracy became significant at around TR 3-4 (6 to 8 seconds after stimulus onset), peaked slightly later, around TR 4-6. Once the distractors became decodable, they remained decodable until the end of the trial. These results suggest that distractor-related information entered all three ROIs, consuming neural resources, which formed the basis for representational competition with the target-related information.

**Figure 3.**
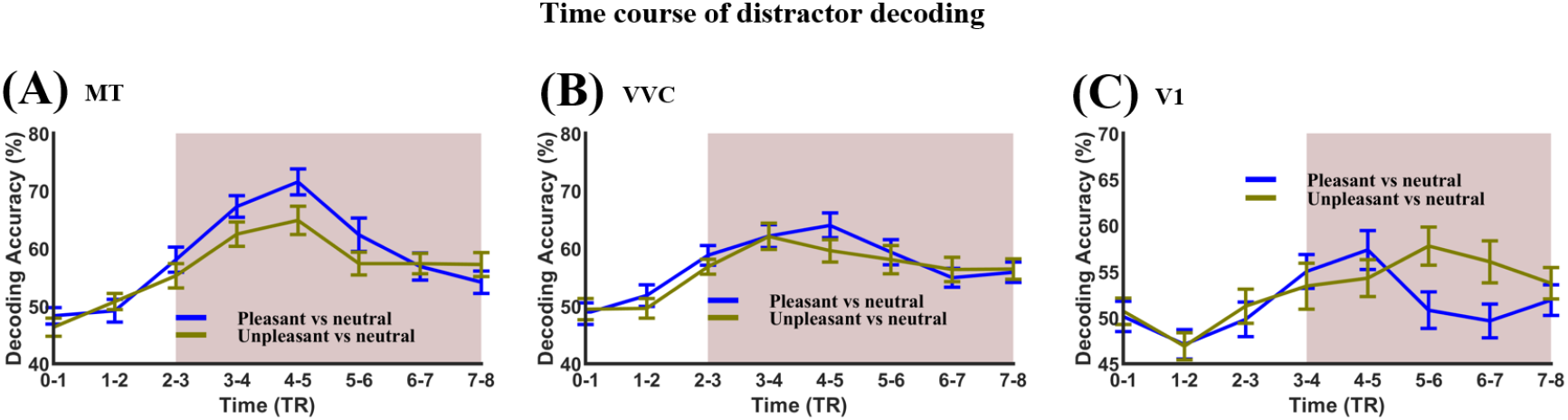
Distractor decoding accuracy time course in MT, VVC, and V1. The shading marks the TRs where the decoding accuracy was significantly higher than chance level (50%). More specifically, in MT (A), the decoding accuracy went above chance level at TR 2-3, in VVC (B), the decoding accuracy went above chance level at TR 2-3, and in V1 (C), the decoding accuracy went above chance level at TR 3-4.

#### Behaviorally-informed decoding analysis

Participants were divided into two groups: high performance group (coherent motion detection accuracy>60%) and low performance group (detection accuracy<60%), and their distractor decoding accuracy time courses were plotted separately and compared in the three ROIs. In MT, Figure 4(A), individuals with lower task performance exhibited a markedly higher distractor decoding accuracy at TR 2-3 for unpleasant vs neutral decoding, suggesting stronger distractor representations at this TR in the low performance group than the high performance group. No significant difference was found between pleasant vs neutral decoding accuracy time courses. In VVC, Figure 4(B), the lower task performance group exhibited a markedly higher distractor decoding accuracy at TR 2-3 for pleasant vs neutral decoding and TR 3-4 for unpleasant vs neutral decoding, respectively, suggesting stronger distractor representations in the low performance group. In V1, Figure 4(C), no significant difference was found in distractor decoding accuracies between the two groups, suggesting that the distractor processing in V1 did not directly impact behavior.

**Figure 4.**
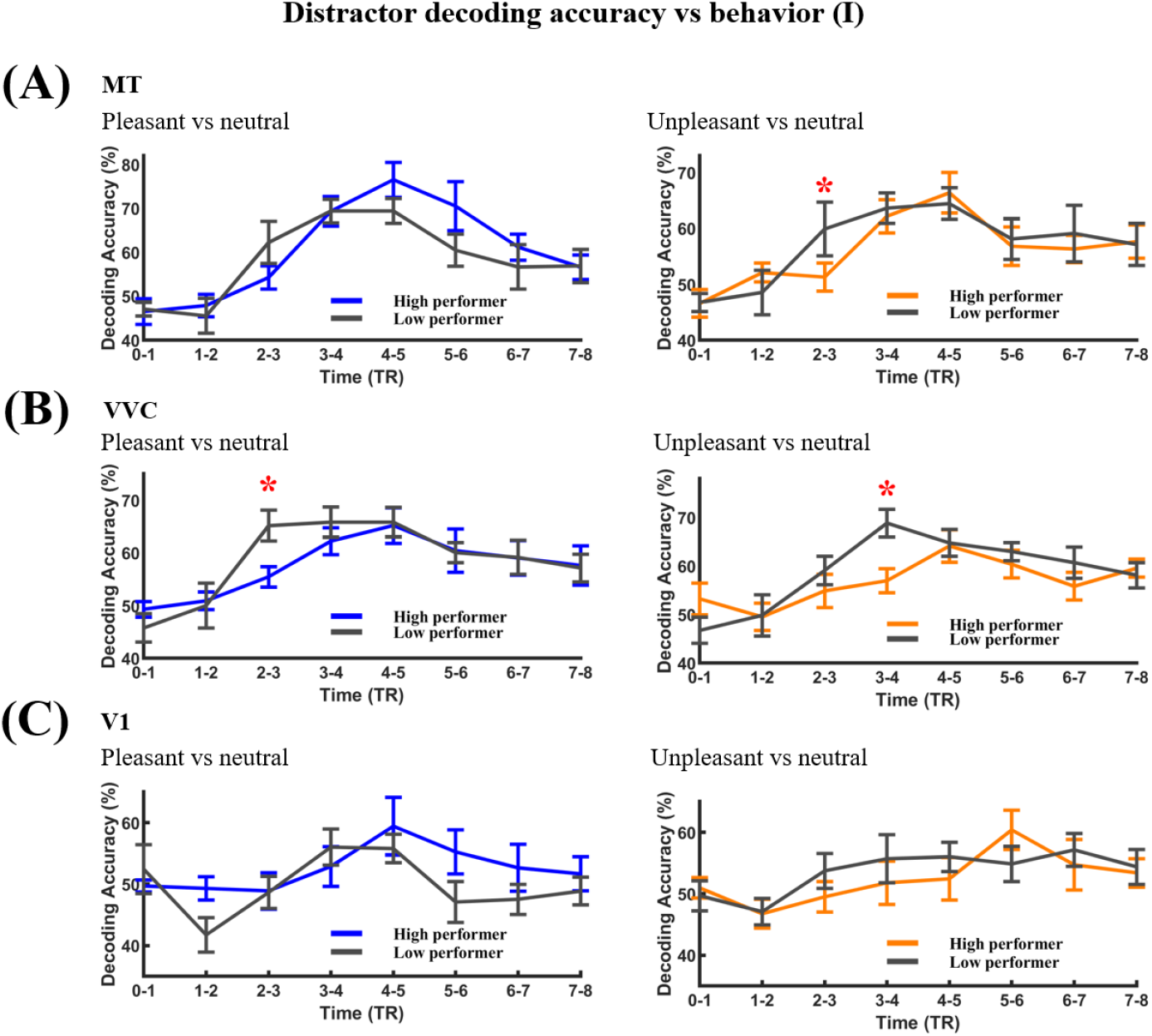
Behaviorally-informed distractor representation analysis I. The subjects were divided into two groups based on their task performance (high performers vs low performers) and the decoding accuracy time courses from the two groups were compared. The red asterisk indicates the TRs where high and low performers show significant difference in distractor decoding accuracy. (A) MT, (B) VVC, and (C) V1.

Next, treating the TRs showing significant differences in the decoding accuracy time courses between high- and low-performing individuals in Figure 4 as TRs of interest, we correlated task performance between decoding accuracy and behavioral performance across participants at all TRs of interest. As shown in Figure 5(A) and 5(B), statistically significant negative correlations were found in VVC for pleasant vs neutral at TR 2-3 (r = −0.6146, p = 0.0018) and for unpleasant vs neutral at TR 3-4 (r = −0.5461, p = 0.0070), and in MT for unpleasant vs neutral at TR 2-3 (r = −0.4262, p = 0.0426), further demonstrating that stronger distractor representations in these motion processing areas were associated with poorer task performance, in support of the hypothesis.

**Figure 5.**
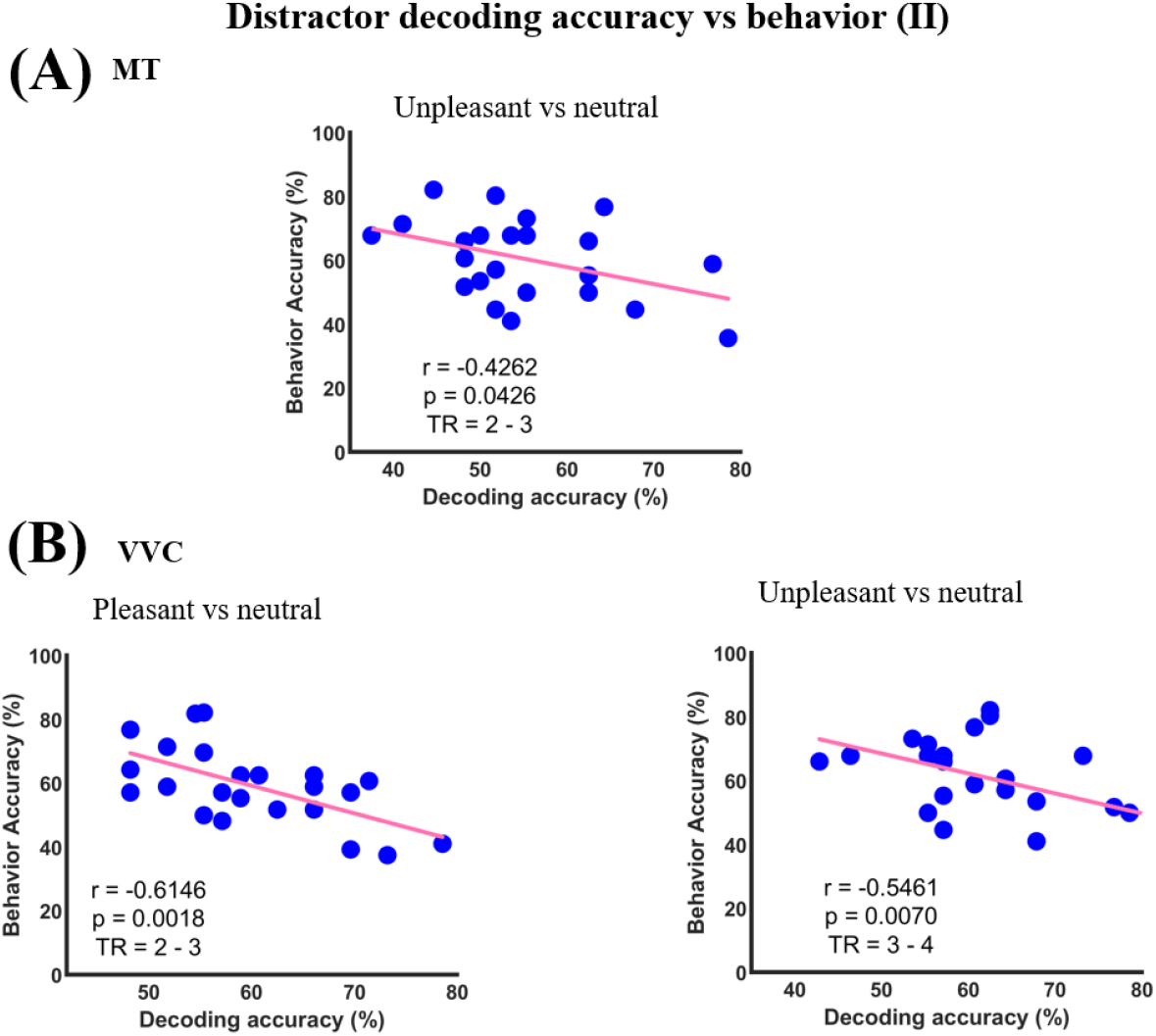
Behaviorally-informed distractor representation analysis II. For all TRs of interest, significant negative correlation across participants was found between decoding accuracy and behavioral performance, suggesting that the stronger the distractor representation, the worse the behavioral performance. (A) MT. (B) VVC.

To quantify the predictive value of visuocortical decoding relative to performance, we built a multiple regression model which used the decoding accuracy values to predict the task performance, pooling the data from both MT and VVC and from all TRs of interest. As shown in Figure 6A, in the model, the detection accuracy of coherent motion was the predicted variable and the decoding accuracies at the TRs of interest across MT and VVC are the predictor variables. For both pleasant vs neutral decoding (Figure 6B) and unpleasant vs neutral decoding (Figure 6C), the predicted coherent motion detection accuracy showed a strong correlation with the true coherent motion detection accuracy, r = 0.6336, p = 0.0012 for pleasant vs neutral and r = 0.6070, p = 0.0011 for unpleasant vs neutral; the model thus was able to explain more than 36% of the variance.

**Figure 6.**
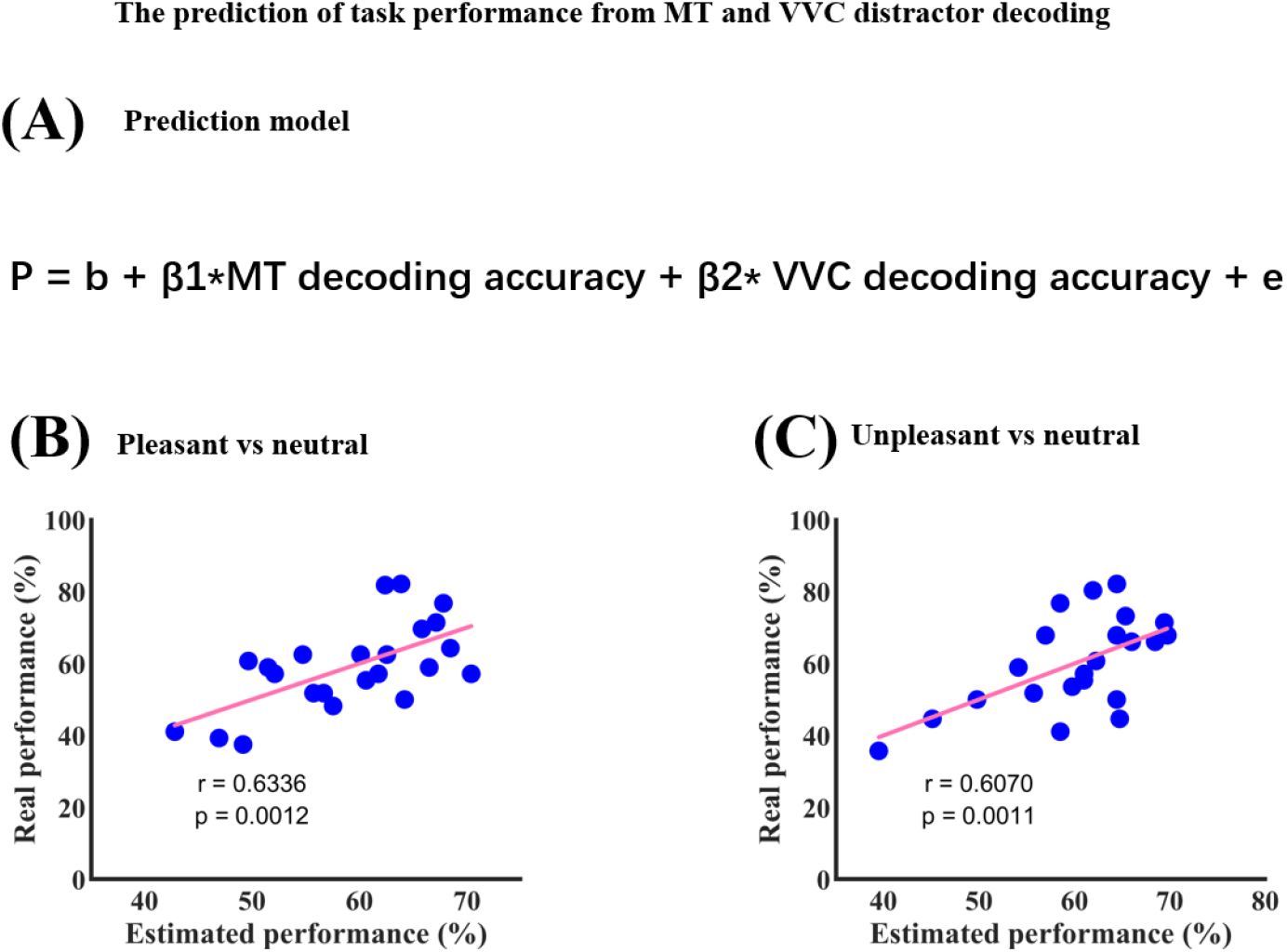
Predicting task performance. (A) Linear model combining MT and VVC decoding accuracies to predict task performance. (B) Pleasant versus neutral. (C) Unpleasant versus neutral.

## Discussion

In real world settings, task-relevant information (targets) and task-irrelevant information (distractors) often appear together, overlapping in space and time. How they compete with each other for neural representations and how the competition impacts behavior is subject to ongoing debate. In this study, we recorded fMRI data from participants performing a motion detection task in which random moving dots (target) flickered at one frequency were superimposed on IAPS pictures (distractor) flickered at another frequency. Participants were instructed to report the number of times the random dots moved coherently. The main results showed that (1) the distractor information was represented in the three ROIs considered, including V1, MT, and VVC, and (2) the stronger the distractor representation in MT and VVC, the worse the performance at detecting coherent motion. These results revealed a competitive relationship between the neural representations of the target and the distractor and the push-pull relationship has a direct impact on behavioral performance.

### Effect of distractors on behavioral performance

In terms of task accuracy, we found that the three types of distractors (pleasant, neutral, and unpleasant) exhibited similar distracting effects on behavior. At first blush, this observation seems to be at variance with studies showing that emotionally arousing stimuli (pleasant and unpleasant pictures for the present case) are processed more deeply and rapidly by the brain (Etkin et al., 2011; Öhman et al., 2001; Pessoa, 2008; Vuilleumier, 2005), which, presumably, should lead to greater interference with the main cognitive task. Findings similar to ours have been reported in the literature. In a visual search experiment with a high-load condition and a low-load condition, the authors found that in the low-load condition (Gupta & Srinivasan, 2015), the emotional distractor exerted stronger distracting influence on task performance than neutral distractors, whereas in the high-load condition, emotional and neutral distractors had similar effects on task performance. They argued that in the high-load condition, a selective inhibition process eliminates the processing advantage of emotional stimuli, and reduces their interference with the primary task to levels comparable to neutral distractors. A similar explanation may apply to the present experiment. In our motion detection task, 175 randomly moving dots had to be monitored to detect coherence in motion, which represented high perceptual load. This was further supported by the relatively low performance with detection rate of coherent motion around ∼60%.

### ROI selection

Three ROIs were considered in this study. MT, located near the dorsal/ventral junction of the superior temporal sulcus, is a hub of the primate motion-processing pathway. Single-unit and lesion work shows that the vast majority of MT neurons are sharply tuned for the direction and speed of moving edges or dot patterns (Albright, 1984; Maunsell & Van Essen, 1983). When local motion signals have to be integrated over space or time, as in the case of a random-dot kinematogram, MT tracks the coherence of global motion and contributes to perceptual decisions on a trial-by-trial basis (Britten et al., 1992; Newsome & Pare, 1988). The VVC—encompassing V4, ventral occipital areas VO1/VO2, and the PHC—also contributes to motion processing. V4 neurons are sensitive to motion-defined boundaries and to the interaction of motion with color/texture cues (Ferrera et al., 1994; Roe et al., 2012). Ventral occipital cortex (VO) responds strongly when motion signals specify three-dimensional form or when objects are defined purely by kinetic boundaries and supports the extraction of motion-defined shapes in the ventral pathway (Dupont, 1997; Van Oostende, 1997). The parahippocampal cortex (PHC) integrates scene layout with large-field optic flow, helping to encode self-motion and navigational context during locomotion (Epstein, 2008; Sherrill et al., 2015). V1, as the first stage of cortical visual processing, receives input from the lateral geniculate and extracts basic visual features such as orientation, spatial frequency, and color, and forwards them to higher-order visual areas for more specialized processing (McManus et al., 2011; Tootell et al., 1998). We included V1 here mainly as a control region to test the specificity of the effect of target-distractor competition on behavioral performance.

### Distractor processing in the ROIs

Although univariate analysis demonstrated robust stimulus-evoked activation in all three ROIs, within the current experimental design, it was not possible to distinguish distractor activation from target activation. The MVPA approach overcomes this limitation. In a previous study, recording fMRI data from participants viewing pleasant, neutral and unpleasant pictures from the IAPS database and applying MVPA decoding to the fMRI data from visual cortex, we showed that pictures from the three emotion categories evoked distinct neural patterns, and the accuracy of decoding one emotion category against another was above-chance in all retinotopic visual areas (Bo et al., 2021). Inspired by this, we divided the trials in the present experiment into the same three emotion categories based on the content of the distracting picture, and performed a similar analysis across TRs. In all three ROIs, the decoding accuracy began to increase as the trial progressed, reaching and staying above chance level at around TR 3, suggesting that the distractor information is processed in the three ROIs in the presence of an attention-demanding primary task. Furthermore, the analysis provided an index, the decoding accuracy, which allowed us to measure the strength of distractor processing at the level of individual subjects, paving the way for evaluating the quantitative relation between distractor processing and behavior.

### Effect of multivariate representational competition on behavior

When task-irrelevant information enters task-relevant brain regions, they compete for neural representations with the primary task. Given limited resources, stronger representations of the distractor imply weaker representations of the target, forming a type of push-pull competitive relationship. For the present study, when the IAPS pictures are better represented, the neural resources for the motion detection task will be reduced, leading to worse motion detection performance. We tested this hypothesis by first dividing the subjects into high and low performer groups and compared their distractor decoding accuracies. In MT and VVC, the low performers show higher distractor decoding accuracy than the high performers at certain TRs, in support of the hypothesis. We then performed regression analysis across participants at those TRs of interest in MT and VVC, to reveal a significant negative correlation between decoding accuracies and task performance, further supporting that participants with stronger distractor representations in these motion processing areas exhibited poorer coherent motion detection performance. Importantly, in V1, no such results were found, demonstrating that the adverse effect of multivariate representational competition on behavior is specific to the visual areas that are engaged both by the primary cognitive task and the distractor.

### Extant literature on spatially and temporally overlapping stimuli

Past studies have considered spatially and temporally overlapping stimuli in the context of how attention reweights their representations in favor of the attended stimulus (Reynolds & Desimone, 2003). At the multivariate representational level, (Kamitani & Tong, 2005) first identified the multivariate pattern evoked by each of the two orthogonally oriented gratings, and then showed the participants the two gratings superimposed and asked them to attend one while ignoring the other. They found that the multivariate pattern evoked by the compound stimulus aligned with the pattern evoked by the attended grating in accord with the prediction of the reweight theory. For spatially and temporally overlapped natural stimuli, faces and houses, category-specific attention reweighting were found in cortical areas FFA and PPA (Cohen & Tong, 2015). Reweighting can take place even without attention manipulation. (Seymour et al., 2009) presented two spatially and temporally overlapping dot fields with different colors and moving directions to the participants. Defining two conjunctions: (red CW + green CCW) versus (red CCW + green CW), they used SVM to predict conjunction pair based on voxel patterns in V1, demonstrating conjunction coding at the earliest cortical stage. Voxel-weight maps, revealing opposite-signed contributions from corresponding voxels for the two conjunctions, are consistent with reweighting patterns. Our study is similar to these studies in the sense that we all considered spatially and temporally stimuli and their multivariate representations in the brain.

The difference lies in that our study asked different questions and, by directly examining the representations of the distractors, we were able to establish the behavioral consequences of these representations despite the fact that attention seeks to minimize these representations.

### Limitations

Our work has a number of limitations. First, the experimental paradigm lacked a no-distractor baseline condition. This leads to our inability to obtain separately, at the univariate level, the contribution from the target and that from the distractor. Second, the target (the moving dots) cannot be grouped into different decodable categories. Consequently, we were not able to examine directly the multivariate representational competition between the target and the distractor at the neural level, and had to, instead, resort to indirect inference by examining the relation between the multivariate representation of the distractor and the behavioral performance. Third, EEG data were recorded along with fMRI in this study. No fusion was attempted between simultaneously recorded fMRI and EEG. However, given that this study concerns specific brain regions in which the target and the distractor compete for representations, it is felt that EEG data, which is known to possess low spatial resolution, had limited potential to contribute.

### Summary

In this work, we reported three main findings: (1) Distractors from different emotion categories exerted similar distracting effects on behavior, (2) distractor information can be decoded in all three ROIs: MT, VVC, and V1, and (3) the strength of the distractor representation indexed by decoding accuracy in MT and VVC, but not in V1, is negatively related with behavioral performance. These results support the notion that a push-pull competition exists at the multivariate representation level between concurrent targets and distractors. In line with the overarching hypothesis of this study, this competition directly impacts task performance.

## Acknowledgements

This work was supported by NSF grants BCS2318886 and BCS2318984 and NIH grant R01 MH125615.

